# Nudibranch predation boosts sponge silicon cycling

**DOI:** 10.1101/2022.08.16.504137

**Authors:** María López-Acosta, Clémence Potel, Morgane Gallinari, Fiz F. Pérez, Aude Leynaert

## Abstract

Sponges are singular players in the marine silicon cycle. They accumulate vast stocks of biogenic silica within their bodies and in the sediments beneath them over long periods. These silica stocks are recycled at slow rates, much slower than that of other silicon users such as diatoms. The observation of an abrupt change in sponge biomass in a temperate coastal ecosystem led us to study the effect of nudibranch (*Doris verrucosa*) predation on the silicon budget of a sponge (*Hymeniacidon perlevis*) population on an annual scale. Predation rates and the associated sponge silicon fluxes were determined. After 5 months of predation, the abundance of sponge individuals did not change but their biomass decreased by 95%, of which 48% can be explained by nudibranch predation. About 97% of sponge spicules ingested by nudibranchs while feeding was excreted, most of them unbroken, implying a high rate of sponge silica deposition in the surrounding sediments. After predation, sponges partially recovered their biomass stocks within 7 months. This involved a rapid growth rate and large consumption of dissolved silicon, with the highest rates ever recorded unexpectedly occurring when the dissolved silicon concentration was minimal in seawater (< 1.5 μM). These findings reveal that the annual sponge predation-recovery cycle triggers unprecedented intra-annual changes in sponge silicon stocks and boosts nutrient cycling. They also highlight the need for intra-annual data collection to understand the dynamics and resilience of sponge ecosystem functioning.

## Introduction

Silicon is a key nutrient in the ocean necessary for a variety of marine organisms to build their siliceous skeletons and grow. These organisms, known as silicifiers, include organisms of different trophic levels and ecological relevance, such as diatoms, silicoflagellates, most species of rhizarians and sponges, and some species of choanoflagellates^1^. Among them, diatoms and sponges are the most abundant pelagic and benthic silicifiers, respectively. Diatoms, which are microscopic unicellular eukaryotic algae, account of about 40% of ocean primary productivity and export of particulate carbon from surface to deep ocean^2,3^. Sponges, which are benthic macroinvertebrates, contribute to the recycling of nutrients to higher trophic levels through benthic-pelagic coupling and to increasing marine biodiversity of micro- and macro-organisms^4–6^. Therefore, the presence of silicon in marine ecosystems maintains the populations of silicifiers, which are key to sustaining the marine biodiversity and food webs that ultimately sustain much of the marine life and human populations^7,8^.

Among silicifiers, sponges are the only group of metazoans (i.e., animals). They are multicellular, heterotrophic organisms with long lifespans, which can last from years to millennia^9,10^. Sponges are common components of marine benthic fauna across global oceans, being numerically abundant and biomass dominant in many marine ecosystems from polar to equatorial latitudes^5,11^. As other silicifiers, sponges consume dissolved silicon from seawater to build their skeletons made of biogenic silica (i.e., opal). These skeletons are more resistant to dissolution than those of other silicifiers, for reasons yet to be determined^12^. Their biological and skeletal characteristics make siliceous sponges major silicon sinks in marine ecosystems, accumulating vast stocks of biogenic silica within their tissues and in the sediments beneath them^12,13^, and recycling their silicon at significantly slower rates (200 to 1000 times slower) than that of other short-living, unicellular silicifiers such as diatoms^14^. Still, in areas where they form large aggregations (e.g., sponge grounds), sponge silicon cycling can enrich in dissolved silicon demersal layers on an oceanographic scale^15^.

The dynamics of sponge populations, which have a major impact on the fate of sponge silicon, strongly depend on recruitment, longevity, substrate competition, and predation^16^. In particular, predation can affect sponge community structure and ecosystem processes^17,18^. Sponge predation has been reported on tropical and subtropical sponge coral reefs^19–22^, mesophotic sponge reefs^23,24^, and at polar latitudes from intertidal to deep-sea environments^25,26^. Spongivory, a term for the condition of feeding on sponges, has been described in some species of mollusks, echinoderms, fishes, and turtles. These organisms may feed exclusively on sponges (i.e, strict spongivores) or have an omnivorous diet that includes sponges (i.e, facultative spongivores). In both cases, sponge predators are adapted to deal with the chemical defenses and mineral skeletons of sponges^27,28^. Among mollusks, dorid nudibranchs (Order Nudibranchia, Family Dorididae) are among the most diverse groups of sponge predators. Most of them are strict spongivores that have adapted their teeth system (radula) to the specific sponge skeletal organization on which they feed^29^. Many of them also have a cryptic coloration with their preys^28^. This high predator-prey specificity implies that most species of dorid nudibranchs feed on one or a couple of sponge species, on which they tend to feed avidly to supply their energetic needs^26,27^.

Monitoring of sponge populations subjected to predation has shown that sponges can largely recover the biomass eaten, at regeneration rates much higher than the growth rate of non-predated sponges^30^. Whether sponge predation and subsequent regeneration processes could affect sponge silicon cycling has never been investigated. In this study we surveyed for one year the population of the sponge species *Hymeniacidon perlevis*, a globally-distributed species^31^ that dominates the sponge community of the Bay of Brest (France), the area of study. In this shallow-water ecosystem, the sponge *H. perlevis* is subjected annually to predation by the nudibranch *Doris verrucosa*. We determined the rates of predation by *D. verrucosa*, which had never been determined. Sponge silicon export during predation and sponge silicon production during regeneration were also determined, and the results were compared with the sponge silicon cycle of non-predated sponges.

## Materials and Methods

### Study species

The sponge *Hymeniacidon perlevis* and the nudibranch *Doris verrucosa* were selected to study the effect of predation on sponge silicon. These species coexist at the maerl bed of Lomergat (48° 17.197’ N; 4° 21.279’ W), a shallow-water ecosystem (depth=3-11 m) located in the Bay of Brest (France). In this ecosystem, which occupies a small area of the Bay (1.44 km^2^ of the total 133.13 km^2^), assemblages of the maerl species *Lithothamnion corallioides* serve as the substrate for a highly diverse benthic community, including more than 20 sponge species^14^. Among them, *H. perlevis* is the dominant species in terms of both abundance (53%) and biomass (64%)^14^. It has a moderate content of siliceous skeleton derived from the production of a single, mid-size (175–475 μm long) type of needle-like silica spicule^32^. This species, which occurs from the intertidal to the subtidal zone in the study area throughout the year, grows on the hard substrate the maerl pieces offer (Fig. 1a).

**Figure 1.**
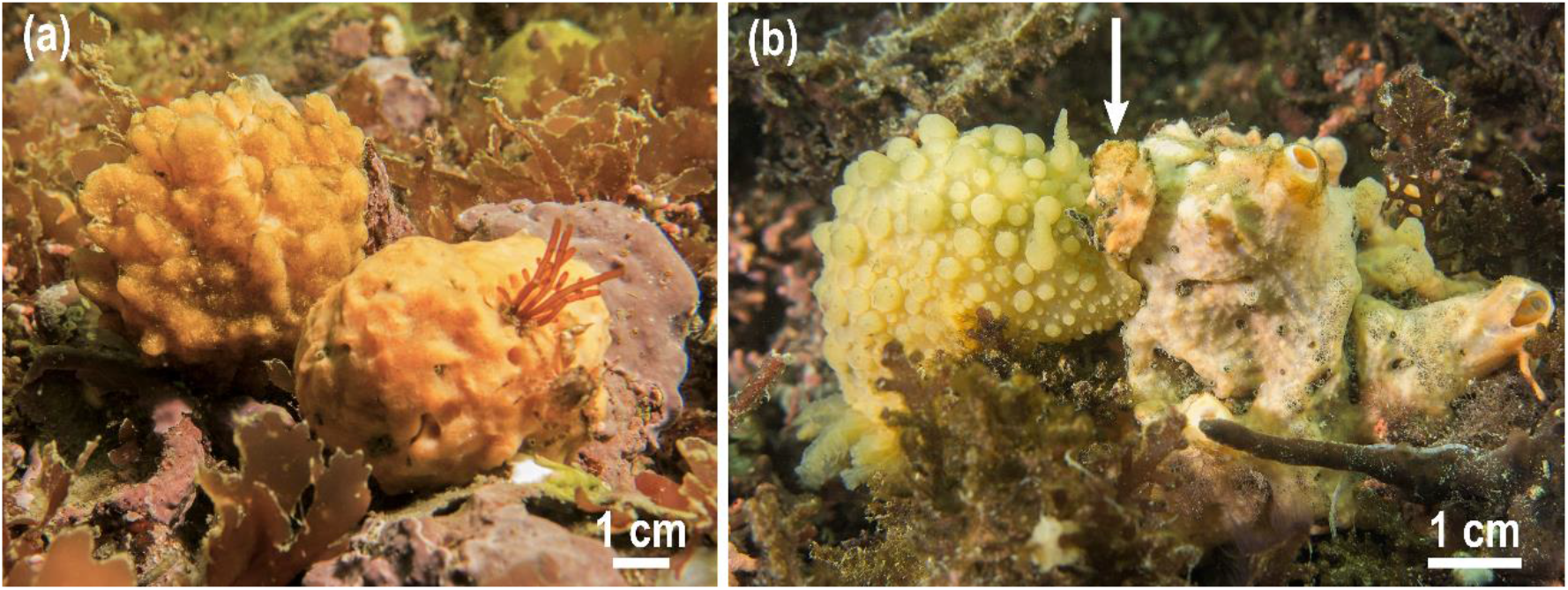
(**a**) View of two individuals of *Hymeniacidon perlevis* growing on the maerl beds of the Bay of Brest. (**b**) An individual of *Doris verrucosa* (left) feeds on the sponge *H. perlevis* (right). The arrow indicates the wound made by the nudibranch when feeding on the sponge.

The area is also habitat to several species of dorid nudibranchs —*D. verrucosa, D. pseudoargus* and *Jonnura tomentosa*— whose adults are present at different seasons of the year: *D. verrucosa* occurs from June to November and *D. pseudoargus* and *J. tomentosa* from October to April. The target species *D. verrucosa* is the dominant one, being at least one order of magnitude more abundant than the other species. The nudibranch *D. verrucosa* lives from the intertidal zone to 15 m depth along the Atlantic coast of Europe and the Mediterranean Sea, and is a strict spongivore^33^. This species has been reported feeding on the sponges *Halichondria panicea*, *H. perlevis* and *Suberites carnosus* in the Spanish Atlantic coast and the Mediterranean Sea^34,35^. These three sponge species occur in the Bay of Brest, but we have only found *D*. *verrucosa* feeding on *H. perlevis* (Fig. 1b).

### Prey abundance

The abundance and biomass of *H. perlevis* in the maerl bed of Lomergat were surveyed for one year (June 2020 – June 2021) by 1×1m random quadrats (n=71) during scuba diving at 4 times over the year: June 2020, to assess the sponge population before predation; November 2020, to evaluate the sponge population after predation; and April and June 2021, to assess the sponge population recovery before the new predation event. During field surveys, each individual of *H. perlevis* found within the quadrats was counted and the linear parameters needed to calculate the volume (length, width, height, diameter) were measured in situ with a ruler. To determine the sponge volume, the body shape of each individual was approximated to one or a sum of several geometric figures (e.g., sphere, cylinder, cube), a non-destructive method widely used in monitoring sponge populations for its well-performance in sponge community assessment and survey^36^. Differences in sponge abundance and biomass between sampling times were examined using one-way Kruskal-Wallis analysis after the normality test (Shapiro-Wilk) failed. When significant differences were reported, post-hoc pairwise multiple comparisons between groups were conducted using the non-parametric Dunn’s test. Statistical analyses —with a significance level of 0.05— and data visualization were performed using the statistical software Sigmaplot 14.5 (Systat Software Inc.).

At each sampling time, 5 individuals of *H. perlevis* were sampled to determine the specific silicon content. Sponge tissue from each individual was sampled (~1-5 mL) and then dried at 60°C to constant dry weight (g). Each dried sample was digested for 5h in 20mL of hydrofluoric acid at 10% (2.9N) as no interference from lithogenic silicon was expected because the samples only contained sponge tissue. After complete digestion, samples in hydrofluoric acid were neutralized with a saturated solution of boric acid (H_3_BO_3_, 60g L^-1^) and dissolved silicon was measured according to the molybdate-blue method by colorimetry on a SEAL Analytical AA3 HR auto-analyzer^37^. Differences in sponge silicon content between sampling times were examined using one-way ANOVA. Data on the specific silicon content and sponge biomass measured in situ after predation (i.e., November 2020, April and June 2021) were used to estimate the rate of sponge silicon production by *H. perlevis* during population recovery when predators were absent (mid-November to mid-June).

### Predator abundance

The abundance and biomass of *D. verrucosa* was studied using 8 transects of 25m x 2m during scuba diving in late June and early July 2020 (total area = 400 m^2^), after the nudibranch population had exploded. Each *D. verrucosa* found within the transects was counted and its size (length) was measured. Whether the nudibranch was feeding on *H. perlevis* or not was also registered.

Nudibranch shape was approximated to that of an ellipsoid (equation 1):

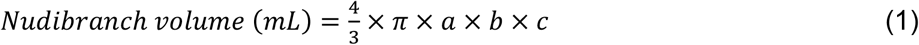

being ‘*a*’, ‘*b*’ and *‘c’* the radius (in cm) of each ellipsoid axes, corresponding to half of the length, width and height of the animal, respectively. These parameters were measured in 36 nudibranchs in the laboratory, together with the wet weight (g). The pairwise relationships between the various morphometric parameters were explored by regression analyses to identify relevant equations that would permit reconstructing biomass reliably from the linear morphometric parameters (length) obtained in the field survey. This allowed us to estimate nudibranch biomass with a non-destructive method. All measured nudibranchs were released into their natural habitat after manipulation.

### In situ nudibranch feeding experiments

Individual predation rates of *D. verrucosa* on *H. perlevis* were determined in situ. To this end, cylinder cages of 20-cm height and 25-cm diameter were built with two meshes that were stitched together with a nylon line (Fig. 2). The smaller net had a 2×2 mm mesh to prevent nudibranchs from escaping. The incubation cages, which were opened on one side, were fixed to the seafloor with steel rebar by scuba diving (Fig. 2a). Twelve incubations were performed in October 2020 in the maerl bed of Lomergat. In nine incubation cages, one individual of *H. perlevis* and one *D. verrucosa* were incubated for 24h (Fig. 2b). Three additional control incubations were performed without a predator. Sponge and nudibranch volumes were measured before and after incubation by measuring the linear parameters in situ with a ruler. Individual predation rate of *D. verrucosa* on *H. perlevis* was calculated as the difference of sponge volume between the beginning and the end of each experiment normalized by nudibranch biomass and time of incubation, following equation (2).

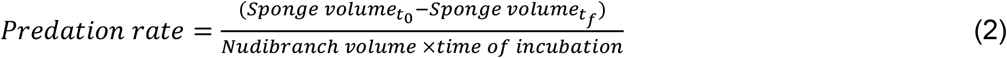

**Figure 2.**
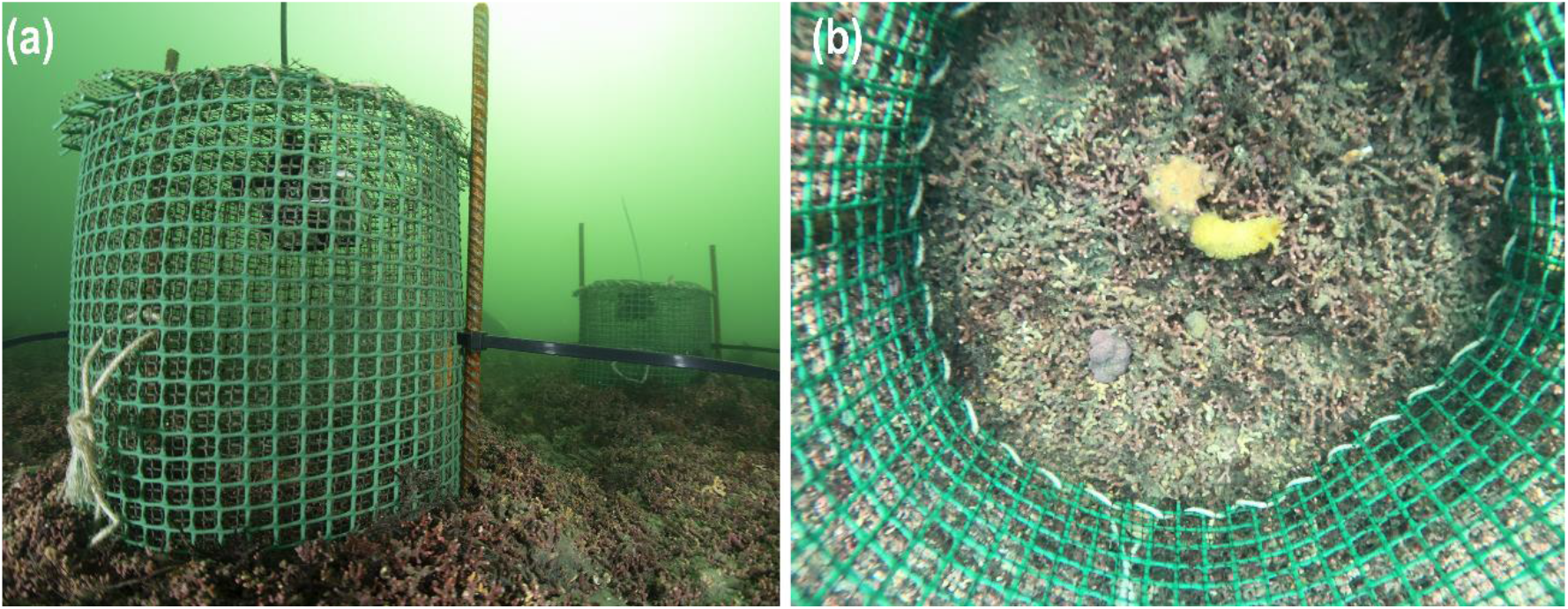
(**a**) In situ experimental setup to determine nudibranch feeding rates on the sponge *Hymeniacidon perlevis* in the maerl bed of Lomergat (Bay of Brest, France). (**b**) Top-down view of the inside of an incubation cage. The nudibranch *Doris verrucosa* approaches to eat the sponge *H. perlevis* after 30 min of incubation.

### Sponge silicon in nudibranch feces

Individual nudibranch feces output was determined in the laboratory on a set of nudibranchs and sponges different from those investigated in situ. Sponges and nudibranchs were acclimated separately before the experiment for 5 days, a period sufficient to allow the nudibranchs to clear their digestive tract from previous feeding^27^. Eight incubations were performed, each incubating a nudibranch with a sponge for 72h in an aquarium filled with running filtered seawater from the Bay of Brest. A 1μm filter pore size was selected to avoid the pass-through of any particle that could eventually settle to the aquaria bottom and be erroneously sampled as nudibranch feces. Acclimation and incubation were performed in an open circuit with a constant flow that ensured the renewal of the water inside each aquarium several times a day.

Feces were sampled with a pipette once a day during incubation and for a 3-day period after the end of the incubation, when fecal production ended. Before further treatment, feces were examined under an optical microscopy with a digital camera (Leica DM2500) to determine their general appearance and the presence of entire or fragmented sponge skeletal pieces (spicules). Samples were then dried at 60°C to constant dry weight (g). To determine the sponge silicon content in nudibranch feces, three subsamples (~8mg) from each incubation aquarium were digested for 24h in 40mL of NaOH 0.2M, a time that ensures the total digestion of sponge silicon in NaOH 0.2M^12^. Dissolved silicon from digestion was measured according to the molybdate-blue method by colorimetry on a SEAL Analytical AA3 HR auto-analyzer^37^. Further, the aluminum content was determined to correct the proportion of lithogenic silica that dissolved at the same time as sponge silicon (i.e., biogenic silica), if any. Dissolved aluminum was determined according to the fluorimetric method using lumogallion^38^. Chemical analysis of aluminum and silicon in the fecal pellets of nudibranchs revealed that the aluminum:silicon ratio was <1.5%, indicating that lithogenic silica in nudibranch feces was negligible. We therefore considered that the silicon measured in the feces was entirely sponge silicon.

To determine the percentage of sponge skeleton ingested that was excreted in nudibranch feces, the amount of sponge eaten by each nudibranch in the aquaria was measured using equation 2, as explained above for the in situ incubations. Also, three samples of sponge tissue (~1mL) of each incubated sponge were sampled to determine the silicon content, which were dried at 60°C to constant dry weight (g) before silicon determination. Silicon content was determined using hydrofluoric acid digestion as for the samples along the sponge field survey (see above). The percentage of ingested sponge skeleton that was excreted in nudibranch feces was determined from the amount of silicon that each nudibranch ingested when feeding and compared with that found in feces.

Individual rates of sponge feeding and fecal production of *D. verrucosa* and the presence of nudibranchs and sponges in the natural habitat were used to determine the amount of sponge silicon deposited on the habitat sediments through nudibranch feces during the predation season.

## Results

### Prey and predator abundance

The field survey revealed that the abundance of *Hymeniacidon perlevis* remained stable throughout the annual cycle, with no significant differences between sampling times (H=3.872, df=3, p=0.276; Fig. 3a). Mean abundance of *H. perlevis* at the study area was 24 ± 17 individuals m^-2^ (n=71). In contrast, sponge biomass significantly changed throughout the annual cycle (H=50.313, df=3, p<0.001; Fig. 3b). The highest sponge biomass value was registered in June 2020 (270 ± 164 mL m^-2^), before the predation season started, and the lowest sponge biomass record was in November 2020 (15 ± 11 mL m^-2^), after the predation season. This implied a loss of sponge biomass from June to November of 95 ± 1 %. Five months after the end of the predation season (April 2021), the sponge biomass was 42 ± 23 mL m^-2^, and 2 months later (June 2021), it was 84 ± 60 mL m^-2^. This means that before the new predation season started, 31 ± 5 % of the initial biomass of *H. perlevis* was recovered (Fig. 3b).

**Figure 3.**
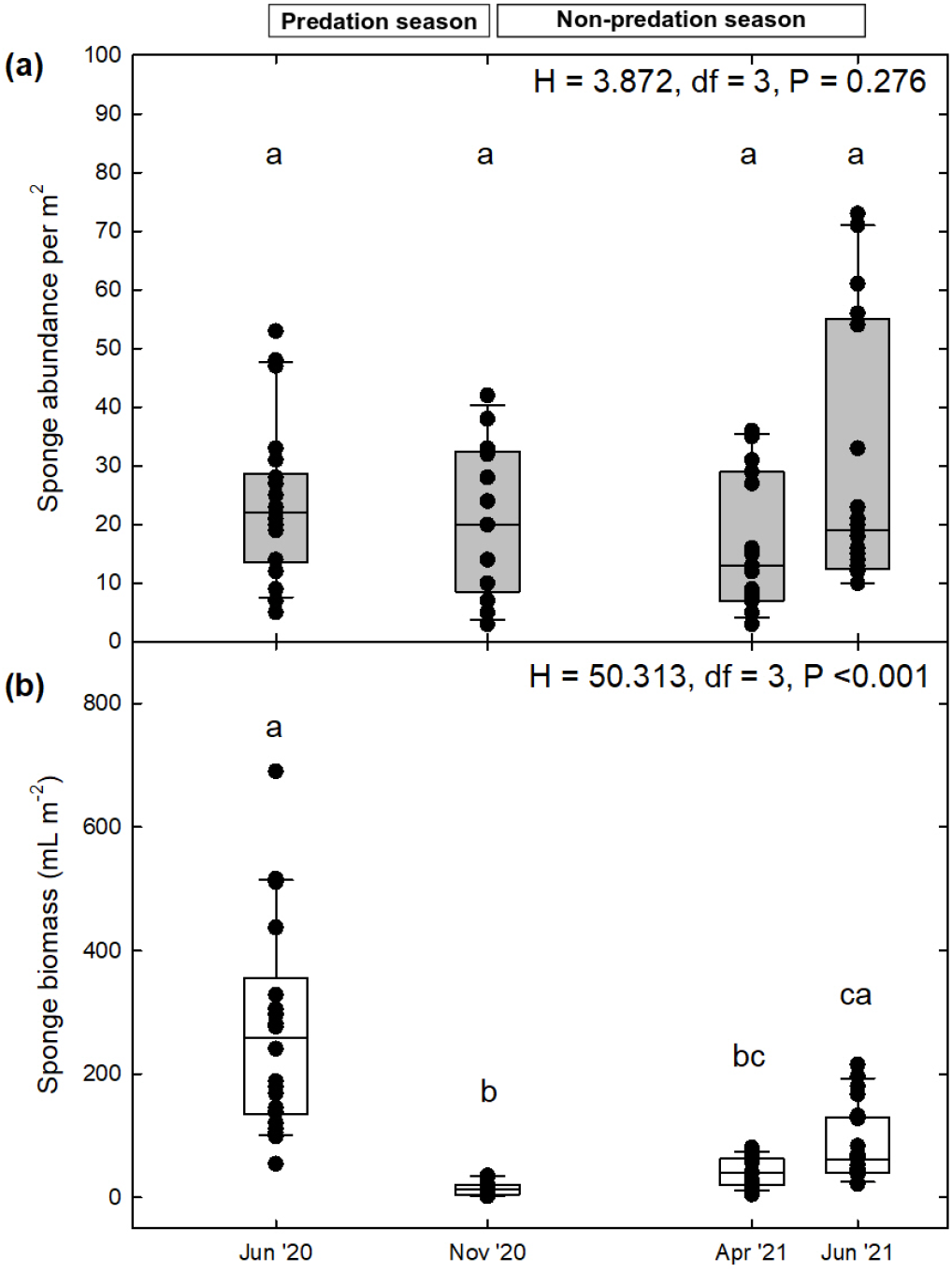
Changes in (a) abundance and (b) biomass of the sponge *Hymeniacidon perlevis* during an annual cycle. Boxplots and individual data points are indicated. Significant between-season differences of (**a**) abundance (individuals m^-2^) and (**b**) biomass (mL m^-2^) of the sponge population of *H. perlevis* are indicated with different letters according to the results of Kruskal-Wallis analysis and the post-hoc pairwise Dunn’s tests.

Nudibranch abundance and size (length) were measured at the beginning of the predation season. A total of 271 individuals of *Doris verrucosa* were registered in 400 m^2^, which were frequently found eating together on a single sponge (up to 8 nudibranchs on the same sponge). Abundance of *D. verrucosa* averaged 0.7 ± 0.4 individuals m^-2^, with a maximum of 9 nudibranchs m^-2^. Nudibranch length ranged from 2 to 8 cm, with a mean length of 4.4 ± 1.3 cm. Individual length values were used to estimate nudibranch biomass through significant correlations between length, volume and wet weight. Relevant linear regressions between nudibranchs length and volume (n=36, R^2^=0.64, p <0.001) and between wet weight and volume (n=36, R^2^=0.82, p <0.001) were established. Average nudibranch biomass in volume was 3.7 ± 2.5 mL m^-2^ and in wet-weight units was 4.5 ± 2.9 g m^-2^. The data of nudibranch volume were used for rate normalization because it facilitates comparison with field measurements that do not require collecting any individual or conducting destructive approaches.

Silicon content of *H. perlevis* per unit of biomass was determined throughout the year and no significant differences between seasons were found (F=0.209, df=3, p=0.888). This indicates that the skeleton content does not vary whether the sponge has been predated or is regenerating its biomass after predation. Mean skeleton content (i.e., biogenic silica) per dry weight of *H. perlevis* was 20.9 ± 8.7%.

Silicon production of *H. perlevis* during regeneration (i.e., mid-November to mid-June) was estimated by combining the species skeletal content and sponge biomass records measured after predation. The biomass of *H. perlevis* increased by 27.7 ± 11.8 mL m^-2^ from mid-November to mid-April, and by 42.0 ± 37.2 mL m^-2^ from mid-April to mid-June, which in terms of silicon is 0.3 ± 0.2 and 0.4 ± 0.5 g Si m^-2^, respectively. Average daily silicon consumption rates were estimated to be 63.7 ± 44.5 μmol Si m^-2^ d^-1^ the first 5 months after predation and 239.1 ± 277.9 μmol Si m^-2^ d^-1^ during spring. All together, silicon production of the sponge *H. perlevis* after predation (i.e., mid-November to mid-June) was 24.2 ± 23.7 mmol Si m^-2^ (0.7 ± 0.7 g Si m^-2^).

### Nudibranch predation rates

The predation activity of *D. verrucosa* was examined in situ. This nudibranch species has only been found feeding on the sponge species *H. perlevis* at the study area. Nine individuals of *D. verrucosa* were incubated for 24h to determine their feeding rate on *H. perlevis*. Individuals of *D. verrucosa* consumed from 0.24 to 1.60 mL of *H. perlevis* per day. Such consumption resulted in a mean normalized predation rate of 0.23 ± 0.11 mL sponge d^-1^ mL^-1^ nudibranch. Sponges incubated without predators (controls; n=3) showed no change in volume.

Individual predation rates measured in situ were extrapolated to the natural habitat, where the abundance of *D. verrucosa* was measured (see above). This resulted in a mean predation rate of 0.84 ± 0.97 mL sponge d^-1^ m^-2^. The large variation associated (i.e., standard deviation) with the predation rate normalized per m^2^ is derived from the patchy spatial distribution of both sponges and nudibranchs. By extrapolating this predation rate to the duration of the predation season (153 days, mid-June to mid-November), the predation rate of *H. perlevis* by *D. verrucosa* was estimated to be 128.0 ± 148.7 mL sponge m^-2^. At this rate, the predation activity of *D. verrucosa* would be responsible for 48 ± 33 % of the sponge population decline measured over the predation season.

### Sponge silicon in nudibranch feces

Fecal production by nudibranchs was examined in the laboratory. The species *D. verrucosa* produced two types of feces when feeding on *H. perlevis* (Fig. 4a-b): colorless fecal pellets, which were unstructured and loaded with sponge spicules (Fig. 4c-d), and colored fecal pellets, which mostly lacked spicules and included organic excretory residues (Fig. 4e-f). Average fecal production was 11.6 ± 8.6 mg DW d^-1^ mL^-1^ nudibranch.

**Figure 4.**
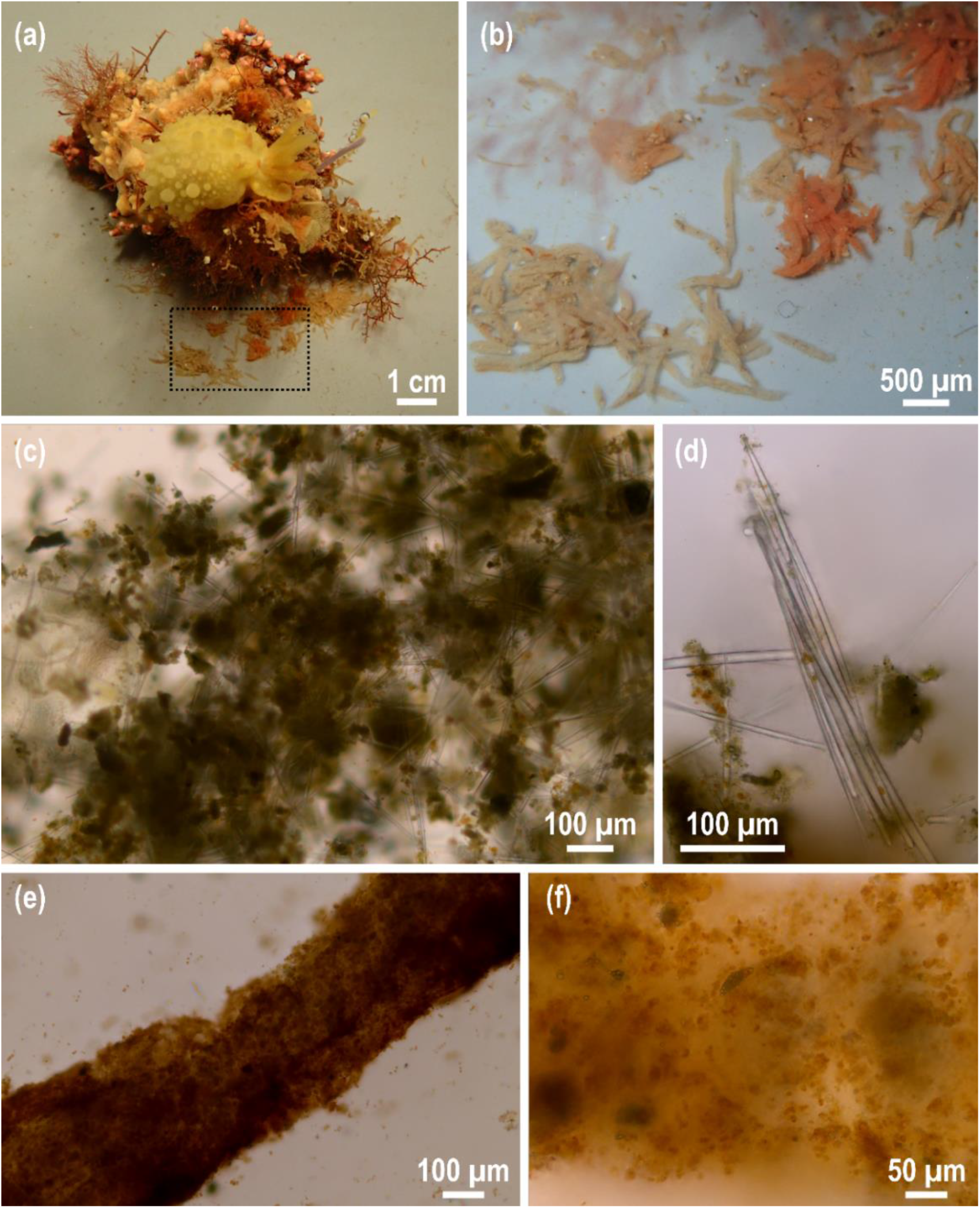
(**a**) The nudibranch *Doris verrucosa* feeds on the sponge *Hymeniacidon perlevis* after 72h of aquarium incubation to assess the silicon export through nudibranch feces. (**b**) Enlargement of the dotted quadrat of Fig. 4a, in which two types of fecal pellets are recognizable: colorless (left) and colored (right) pellets, the latter ones colored with the same pigments existing in the eaten sponge. (**c-d**) Microscopic view of colorless fecal pellets of *D. verrucosa*. Colorless feces, less structured than colored ones, are loaded with sponge spicules, which are mostly unbroken. In some cases (**d**), spicules still have the same structural organization as in the sponge. (**e-f**) Microscopic view of colored fecal pellets of *D. verrucosa*. Colored feces are more structured than colorless ones and mainly contain organic excretory debris.

Silicon content in nudibranch feces ranged from 0.10 to 0.18 g Si g^-1^ DW, and the normalized rate of excretion ranged from 0.3 to 4.3 mg Si d^-1^ mL^-1^ nudibranch. In parallel, the amount of silicon ingested by each nudibranch was estimated by determining the amount of sponge eaten per nudibranch and the silicon content of each sponge examined. During laboratory experiments, nudibranchs ingested 0.4 to 4.1 mg Si d^-1^ mL^-1^ nudibranch. Note that this ingestion rate cannot be compared to that measured in situ, as assayed nudibranchs in the laboratory were starved for 5 days before incubation. When comparing the amount of silicon ingested by each nudibranch with that found in feces, about 97.0 ± 8.7% of the ingested sponge silicon was excreted in nudibranch feces.

Based on the abundance of nudibranchs, the nudibranch predation rate and the specific silicon content of *H. perlevis*, the rate of sponge silicon deposition rate during predation was calculated. Nudibranchs ingested 1.2 ± 1.7 g Si m^-2^ during the predation season (153 days, mid-June to mid-November). Using the ratio of sponge silicon excreted in feces (97.0 ± 8.7%), the rate of sponge silicon deposition through nudibranch feces during the predation season was estimated in 1.2 ± 1.8 g Si m^-2^.

## Discussion

The sponge population survey of *Hymeniacidon perlevis* at the maerl bed of Lomergat (Bay of Brest, France) revealed unexpected changes over the course of a year. From June to November, the biomass of *H. perlevis* drastically decreased in 95 ± 1% (Fig. 3). That period coincides with the season when the nudibranch *Doris verrucosa* is present and feeds on the sponge *H. perlevis*, its only source of food reported in the area of study. At a feeding rate of 0.23 ± 0.11 mL sponge d^-1^ mL^-1^ nudibranch, the species *D. verrucosa* is responsible for 48 ± 33% of the sponge biomass decline. The rest of the sponge biomass lost could be due to the feeding activity of other sponge feeders such as the polychaete *Euphrosine foliosa* and the gastropod *Emarginula fissure*, which have been reported feeding on sponges in the Bay of Brest^39^.

During the predation season, deposition of sponge silicon in marine sediments increases as nudibranchs excrete most of the siliceous skeletons of the eaten sponges in the feces (97.0%; Fig. 4). This result is in agreement with other studies on strict spongivores such as hawksbill turtles, angelfishes, sea stars and other species of nudibranchs, in which high sponge silicon content in digestive tracts and feces have also been reported (68.3 – 99.9%)^19,20,27,40^. The increase of sponge silicon deposition will probably be more locally significant in areas where low-mobility spongivores, such as nudibranchs and sea stars, are present. The high deposition of sponge silicon would in fact enhances the role of sponges as silicon sinks. However, there is also the question of whether once sponge silica (opal) passes through the digestive tract of predators, it may become more fragile to dissolution, which, if so, would imply a feedback mechanism that could help sponges regenerate their biomass eaten.

Sponges usually recover their wounded tissue during the first month after damage, but when the damage is severe, the process can take several months^30,41^. In the study area, the number of sponges did not change during the year (Fig. 3a), but the average individual size decreased from 11.3 ± 20.3 to 0.7 ± 1.5 mL individual^-1^ from June to November, regardless of whether the decrease was due to the predation of *D. verrucosa* or another cause. After such severe biomass decrease, the sponge population recovered a total biomass of 69.7 ± 49.0 mL m^-2^, of which 27.7 ± 11.8 mL m^-2^ were from November to April and 42.0 ± 37.2 mL m^-2^ from April to June (Fig. 3b). The highest sponge growth (regeneration) measured in our study occurred when the availability of the main sponge food (pico- and nanoplankton, including bacteria) was at its highest, that is, from mid-April to June, when plankton blooms occurred^42^. Regeneration in sponges is in contrast with the generally slow growth of these organisms^43^. Regeneration rates can be 2 to 10 times faster than growth rates^41,44,45^, and in some cases even up to 2900 times the natural growth rate^46^. Such high rates not only occur during regeneration, but have also been measured in episodes of high availability of food, such as that following the collapse of part of the ice-sheet Larsen A in the Antarctica, where an impressive sponge population explosion was reported^47^.

To regenerate their damaged biomass, sponges must consume not only food to supply their energetic and growth needs but also dissolved silicon to build their siliceous skeletons as they grow. Unexpectedly, the highest silicon consumption occurred in the time of the year when dissolved silicon in seawater is minimum in the Bay of Brest (April-June; <1.5 μM Si), when diatoms bloom and deplete dissolved silicon availability^48^. Furthermore, the reported sponge silicon consumption from mid-April to mid-June during regeneration would not be fulfilled even if the concentration of dissolved silicon in seawater had been that at which the highest silicon consumption in *H. perlevis* was determined (100-150 μM Si)^49,50^, a concentration never measured in the Bay of Brest and only available in some remote zones such as the Southern Ocean and deep North Pacific Ocean^51^. In a hypothetical exercise, if *H. perlevis* would consume dissolved silicon at its highest silicon consumption rate (V_max_ = 0.134 ± 0.018 μmol Si h^-1^ mL^-1^ sponge)^49^ consumption from mid-April to mid-June, sponge growth would be 31.9 ± 12.1 mL m^-2^. However, the sponge growth measured in the field at that period was 42.0 ± 37.2 mL m^-2^. The molecular and physiological mechanisms behind this high consumption of silicon are enigmatic. Silicon transport in sponges is partially due to passive transporters (aquaglyceroporins), which require a high dissolved silicon concentration outside the sponge to transport silicon within the sponge tissue, where silicon continues its transport and transformation for siliceous biomineralization of the sponge skeleton^52^. Our results suggest that unknown molecular and physiological mechanisms may be unlocked during regeneration to meet silicon requirements during sponge tissue regeneration.

Sponge silicon fluxes of non-predated sponges were estimated to compare with those measured in this study. Silicon production in non-predated sponges was estimated from the species silicon consumption kinetics^49^ at the concentration of dissolved silicon in the bottom waters of the Bay of Brest, which is weekly measured by the SOMLIT-Lanvéoc Observation Service, over the period of study (i.e., June 2020-November 2020 and November 2020-June 2021). It resulted in 0.3 ± 0.1 and 0.7 ± 0.3 g Si m^-2^ for the predation and recovery season, respectively. Silicon deposition in non-predated sponges was estimated from the average sponge silicon deposition measured in the maerl beds of the Bay of Brest (0.9 ± 0.2 g Si m^-2^ y^-1^)^14^ and by considering that the contribution of sponge silicon of *H. perlevis* should be proportional to the contribution of the species to the silicon standing stock in the sponge community per m^2^ of the maerl bed of Lomergat, which is 65.5 ± 21.0 %. It resulted in 0.2 ± 0.1 and 0.3 ± 0.2 g Si m^-2^ for the predation and recovery season, respectively. Hypothetical sponge silicon stock of non-predated sponges was calculated by adding the sponge silicon production and subtracting sponge silicon deposition to the initial silicon stock at each period of study (June 2020-November 2020 and November 2020-June 2021). To compare sponge silicon fluxes of predated and non-predated sponges, the stock-specific flux rates were calculated by normalizing the flux rate (either for production or deposition) to the corresponding sponge silicon stock at the beginning of the season (i.e., predation or recovery). The rate of stock-specific sponge silicon deposition over the predation season is 5 (0.5/0.1) times faster than that estimated for non-predated sponges (Fig. 5), indicating that nudibranch predation boosts sponge silicon transport from the living community to the sediments below. Similarly, the rate of stock-specific sponge silicon production over the recovery season is 16 (4.8/0.3) times faster than that estimated for non-predated sponges (Fig. 5). This indicates that regenerating sponges consume silicon at much faster rates than undamaged sponges to recover their eaten biomass, mobilizing large amounts of silicon. Therefore, in a sponge population subjected to predation, sponge silicon deposition is the dominant flux during predation and sponge silicon production is the dominant flux during recovery (Fig. 5).

**Figure 5.**
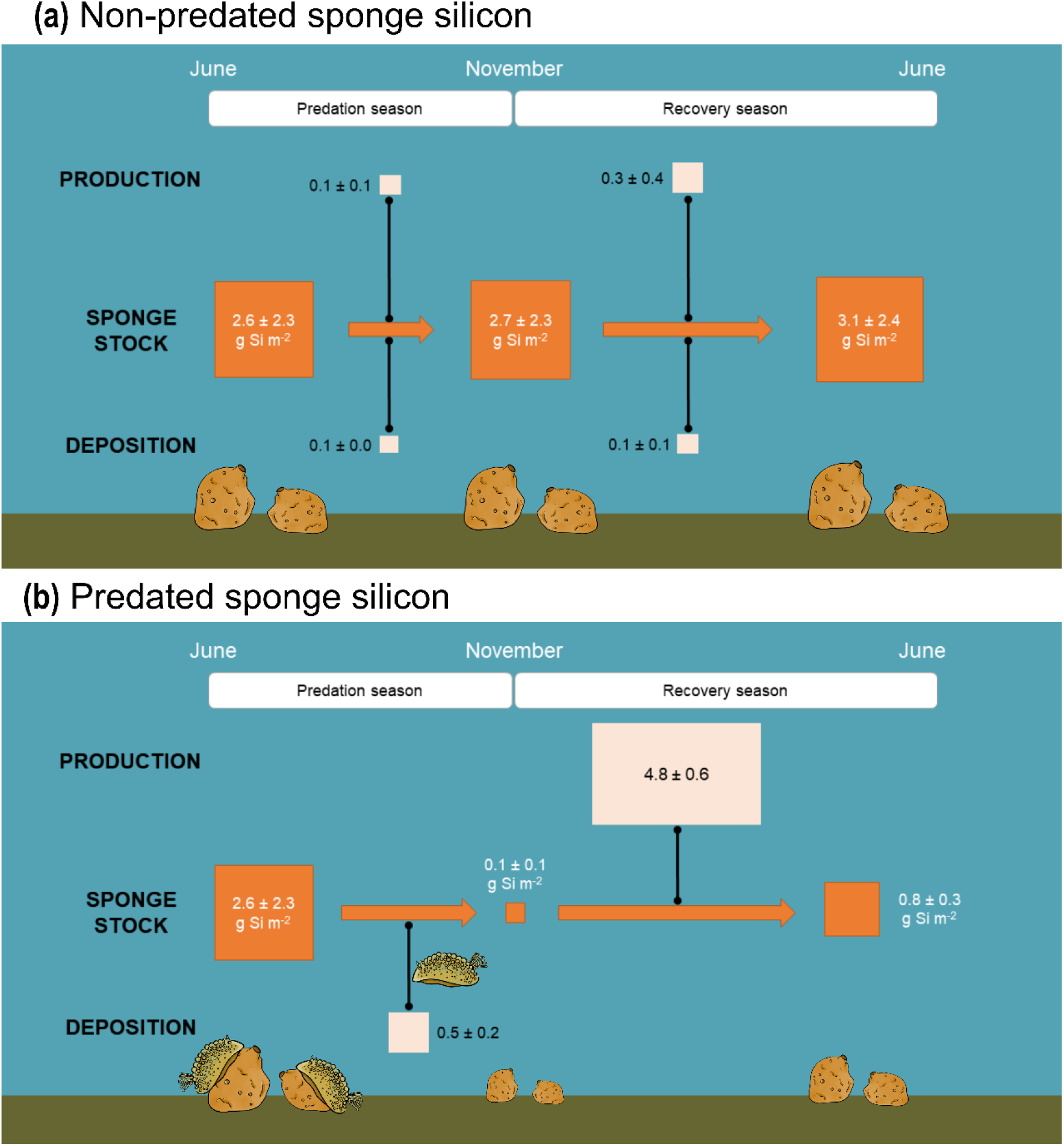
Scheme summarizing sponge silicon (Si) dynamics (**a**) as previously thought when predation is not considered and (**b**) integrating the predation effects measured in this study. Sponge silicon stocks are indicated in g Si m^-2^ and stock-specific flux rates are dimensionless (g Si m^-2^ produced or deposited over the season considered: g Si m^-2^ initial sponge stock). The size of the boxes representing both stocks and stock-specific flux rates is proportional to their value. More details about the calculations are available in the main text.

The changes in sponge biomass measured in this study revealed the importance of intra- and inter-annual surveys for biogeochemical and ecological purposes. In the study area, a punctual sampling to determine the annual sponge silicon budget would make errors of up to three orders of magnitude in both sponge silicon fluxes and stocks, depending on whether the sampling would be done in June or November, for instance, and could omit sponge silicon fluxes that occur primarily in a given period of the year. In other regions such as the North Atlantic Ocean and the Southern Ocean, sponge biomass variations of 40 to 80% have also been attributed to predation and regeneration cycles over months to years^25,53^. This has implications for sponge monitoring planning, especially in the current scenario of anthropogenic climate change, in which marine ecosystems are changing rapidly and marine life is affected. Populations near the limit range of the species geographical distribution or under changing environmental conditions (e.g., ocean warming) can destabilize the predator-prey equilibrium^54,55^. An example of such a disruption has occurred on the coast of Alaska, where a heavy recruitment episode of the spongivore nudibranch *Archidoris montereyensis* led to the disappearance of the sponge species *Halichondria panicea* over an area of 550 square meters^56^. Such loss of the sponge *H. panicea* led to a shift in the dominant species in the area, which is now occupied by the brown alga *Alaria marginata*^26^.

## Conclusion

Our results indicate that sponge predation not only modifies the structure and dynamics of sponge communities but also sponge-mediated biogeochemical cycling. They also show that the intra-annual variations of predated sponge populations need to be taken into account when measuring their silicon budget. Otherwise, biogeochemical estimates disconnected from the ecological dynamics can generate large errors in estimates of nutrient utilization. Our findings revealed unprecedented intra-annual changes in sponge silicon biomass, which are associated with higher rates of sponge silicon utilization than expected, irrespective of the dissolved silicon availability. This cycle of sponge predation-recovery at an annual rate suggests an enhanced nutrient cycling and has important implications for understanding the ecosystem functioning and the benthic-pelagic coupling of nutrients.

## Acknowledgements

The authors thank Thierry Le Bec and Erwan Amice for assistance during underwater fieldwork and for pictures of Fig. 1 and 2a. The authors also thank Eric Dabas for his technical support at the aquarium facilities of the European Institute for Marine Studies and the staff maintaining the SOMLIT-Lanvéoc database for making public their information on nutrient availability at the bottom waters of the Bay of Brest. Natalia Llopis Monferrer is also thanked for her help with the artwork in Fig. 5 and comments on an early version of the manuscript. This research was supported by the French National research program EC2CO (grant 12735 – AO2020) and the ISblue project, Interdisciplinary graduate school for the blue planet (ANR-17-EURE-0015), co-funded by a grant from the French government under the program “Investissements d’Avenir” (grant ARISE - Thème 4 Recherche). MLA thanks the Xunta de Galicia for her postdoctoral grant (IN606B-2019/002), which also supported this work.

## Author contributions

Conceptualization and validation: MLA; Investigation: MLA, CP, MG; Formal analysis: MLA, CP; Visualization: MLA, AL; Funding acquisition, project administration and supervision: MLA, FFP, AL; Writing original draft: MLA; Review and editing manuscript: all authors.

